# Analyzing single-cell bisulfite sequencing data with *MethSCAn*

**DOI:** 10.1101/2022.06.15.496318

**Authors:** Lukas P. M. Kremer, Martina M. Braun, Svetlana Ovchinnikova, Leonie Küchenhoff, Santiago Cerrizuela, Ana Martin-Villalba, Simon Anders

**Affiliations:** BioQuant Centre, University of Heidelberg, Heidelberg, Germany; Division of Molecular Neurobiology, German Cancer Research Center (DKFZ), Heidelberg, Germany

## Abstract

Single-cell bisulfite sequencing (scBS) is a technique that enables the assessment of DNA methylation at single-base pair and single-cell resolution. The analysis of large datasets obtained from scBS requires preprocessing to reduce data size, improve signal-to-noise ratio, and provide interpretability. Typically, this is achieved by dividing the genome into large tiles and averaging the methylation signals within each tile.

Here, we demonstrate that this coarse-graining approach can lead to signal dilution. As an alternative, we propose improved strategies to identify more informative regions for methylation quantification, and a more accurate quantitation method than simple averaging. Our approach enables better discrimination of cell types and other features of interest and reduces the need for large numbers of cells. We also present an approach to detect differentially methylated regions (DMRs) between groups of cells, and demonstrate its ability to identify biologically meaningful regions that are associated with genes involved in the core functions of specific cell types.

To facilitate the analysis of scBS data, we have developed a software tool called *Meth-SCAn* that implements these methods and provides additional functionality.

## Introduction

Sequencing-based assays with single-cell resolution have offered new means to understand the differences between the cells making up a sample. Single-cell RNA sequencing (scRNA-seq) techniques have matured at great pace in recent years, with well developed analysis methodology, and methods to study epigenetics at single-cell resolution are rapidly catching up.

Here, we discuss bisulfite sequencing, a method to study DNA methylation. In brief, DNA is treated with bisulfite, which converts unmethylated cytosines to uracils which are read as thymine in subsequent PCR, while methylated cytosines are protected from conversion. After sequencing, these conversions allow for the determination of the methylation status of all cytosines covered by reads (Frommer et al., 1992). Bisulfite sequencing can also be performed at single-cell resolution (Smallwood et al., 2014) and even in parallel with scRNA-seq (Hu et al., 2016; Clark et al., 2018).

In the present paper, we discuss strategies to analyze single-cell bisulfite-sequencing (scBS) data. We suggest improvements to current approaches, and demonstrate their value in benchmarks, using four real-world datasets. Furthermore, we discuss how to perform comparative analyses. Finally, we present *MethSCAn*, a comprehensive software toolkit to perform scBS data analysis, using the described methods.

### Standard approach

The standard approach to analyze scBS data is based on methodology developed for the analysis of scRNA-seq data. Therefore, we start by briefly reviewing how sc*RNA*-seq data is commonly analyzed, before we discuss scBS data analysis.

The starting point in most scRNA-seq analyses is a matrix of UMI counts (i.e. counts of distinct RNA molecules), one row for each cell and one column for each gene. A first goal is usually to assign cell types or states to cells. To this end, one needs to establish which cells are similar to each other, i.e., quantify the distance (i.e., dissimilarity) between any two given cells’ transcriptional profile. A standard approach, used with minor variation in virtually all recent research and automated by popular software such as Seurat (Hao et al., 2021) or Scanpy (Wolf et al., 2018), is as follows: One first accounts for cell-to-cell variation in sequencing depth by dividing each UMI count by the respective cell’s total UMI count, then transforms to a homoskedastic scale by taking the logarithm. In order to avoid matrix elements with zero count to be transformed to minus infinity, one typically adds a very small “pseudocount” (often 10^−4^) to the normalized fractions before taking the logarithm. Now, one could use Euclidean distances of these vectors of logarithmized fractions as dissimilarity score. However, these scores would be exceedingly noisy due to the strong Poisson noise introduced by the many genes with very low counts. Therefore, one performs a principal component analysis (PCA), keeping only the top few (typically, 20 to 50) components. As Poisson noise is uncorrelated between genes, it will average out in the top principal components, as these are all linear combinations with weight on a large number of genes. Therefore, Euclidean distances between these “PCA space” vectors provide a robust dissimilarity score. Hence, the PCA space representation is suitable as input to methods like t-SNE and UMAP, which provide a two-dimensional representation of the data, or to methods for clustering (assigning cells to groups by similarity) and trajectory finding (identifying elongated manifolds in PCA space and assigning cells to quasi-1D positions along them).

This procedure is commonly adapted when working with single-cell DNA methylation data, because once one gets to the PCA step, one can then continue with the established methods just mentioned. However, constructing a matrix suitable as input for PCA from methylation data requires deviation from the standard scRNA-seq workflow due to considerable differences in data structure. First, while scRNA-seq quantifies RNA abundance of genes or transcripts, scBS is genome-wide and thus lacks a natural choice for features in which methylation is to be quantified. Second, instead of counts, scBS generates binary data that informs us whether certain cytosines in a given cell are methylated or not. A simple and common approach to construct a methylation matrix suitable for PCA, used for instance by Luo et al. (2017), is to divide the genome into tiles of e.g., 100 kb size, and calculate for each cell the average methylation of the DNA within each tile. To this end, one identifies in the tile all CpG sites that are covered by at least one read and averages their methylation state, i.e., one denotes as average DNA methylation of the tile in a given cell the proportion of the observed CpG sites in the tile that were found to be methylated (Fig. 1A). This yields a matrix, with one row for each cell and one column for each genomic tile, comprising numbers (“methylation fractions”) between 0 and 1. This matrix is now subjected to PCA. After PCA, one can proceed with dimensionality reduction and clustering approaches known from scRNA-seq.

**Figure 1.**
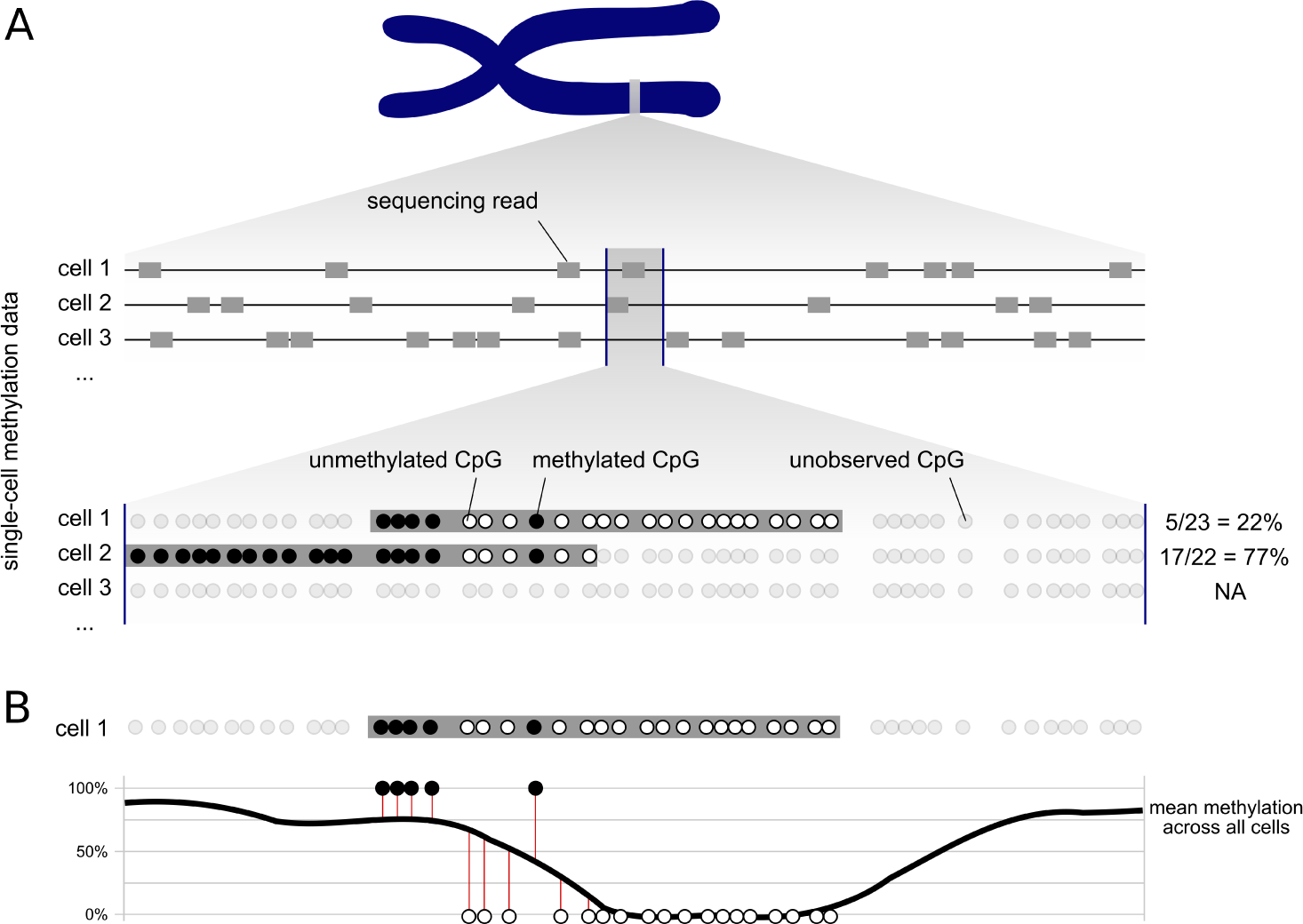
Improved quantification of DNA methylation in a given genomic interval. **(A)** Depicted is a genomic interval (between vertical blue lines) along a chromosome, for which DNA methylation is to be quantified. Two cells cover differing parts of the interval with one read each. If one simply counts for each cell which fraction of the covered CpG sites are methylated, one obtains very different values for the two cells. **(B)** By averaging each CpG site’s methylation over all cells and subsequent smoothing, the thick black “average methylation curve” is obtained. To quantify the methylation of cell 1 from (A) relative to this average over all cells, we propose to use the cell’s residuals to the smoothed curve (lengths of the vertical red lines) and take their average, counting residuals of methylated CpGs positive and residuals of unmethylated CpGs negative.

While this simple procedure is straight-forward and produces usable results, it is not optimal. In this paper, we discuss weaknesses of the simple approach and suggest several refinements to overcome them. Using benchmarks and application to real data, we show that our improvements substantially increase the information content of the processed data. In the main text, we explain the proposed methods and their motivation in a qualitative manner, while mathematical details are given later in the Methods section. Then, we will demonstrate the value of our methods using benchmarks and application to real data. Finally, we describe and demonstrate an approach to detect differentially methylated regions (DMRs) in scBS data. We also describe our software toolkit, *MethSCAn*, that facilitates all these analyses.

## Improved preprocessing of scBS data

### Read-position aware quantitation

We first discuss the task of quantifying the level of methylation in a given, fixed, genomic interval. Typically, read coverage per cell is sparse in scBS data. In the example of Figure 1A, the depicted interval is covered by a single read for two of the three cells shown and no read in the third. The read from cell 2 shows much more methylation than the read from cell 1, and the standard analysis would therefore consider cell 2 to have higher methylation in the interval than cell 1. However, given that the two reads agree wherever they overlap, a more parsimonious interpretation would be that the cells do not show difference in methylation within the interval. Rather, both cells, and similarly maybe most other cells, might have stronger methylation in the left third of the interval than in the middle one.

Therefore, we propose to first obtain, for each CpG position, a smoothed average of the methylation across all cells, and then quantify each cell’s deviation from this average. In Figure 1B, the curved line depicts such an average over all cells, and the red vertical lines show an individual cell’s deviation from the ensemble average. We take the lengths of the red lines as signed values (“residuals”), positive for lines extending upwards from the curve (methylated CpG) and negative for lines extending downwards (unmethylated CpG). For each cell, we then take the average over the residuals for all the CpGs in the interval that are covered by reads from this cell. In this average, we perform shrinkage towards zero via a pseudocount (to trade bias for variance, see Methods for detail) in order to dampen the signal in cells with low coverage of the interval.

The average thus obtained, the shrunken mean of the residuals, is what we use to quantify the cell’s (relative) methylation in the interval. For a genome tiled into such intervals, we thus obtain a matrix, one row per cell, one column per interval, that can be used for downstream analysis, e.g., as input for PCA. The signal-to-noise ratio in this matrix will be better than in a matrix obtained by simply averaging absolute methylation (0 or 1) over all the cells’ covered CpG sites in a region. The reason for this is that we reduce the variation in situations as the one depicted in Figure 1A, where the methylation of the reads might differ strongly even though there is no actual evidence for a difference between the two cells.

How should one obtain the ensemble average (the curved line in Fig. 1B)? A simple approach to get a value for a specific CpG would be to take all cells with read coverage for the CpG and use the fraction of these that show the CpG as methylated. However, especially when only few cells offer coverage, these averages will be very noisy. Therefore, we propose to smoothen using a kernel smoother, i.e., by performing a kernel-weighted average over the CpG site’s neighborhood. The kernel bandwidth (i.e., the size of the neighborhood to average over) is a tuning parameter; for the examples presented here we used 1000 bp.

A minor remaining issue is how to deal with the case that a cell has no reads at all within a given interval. Here, it is justified to simply put zero into the matrix element, because a shrunken residual average of zero indicates that there is no evidence of the cell deviating from the mean. We slightly refine this by using an iterative imputation within the PCA (“iterative PCA”, see Methods for details).

Taken together, this shrunken mean of residuals quantitation reduces variance in comparison to simple averaging of raw methylation calls. We will show further below how this improves results.

### Finding variably methylated regions

Typically, some regions in a chromosome will have very similar methylation status in all cells, while other regions show variability in methylation across cells. For instance, it is known since long that CpG-rich promoters of housekeeping genes are unmethylated, and that a large proportion of the remaining genome is highly methylated regardless of cell type (Bird, 1986). In contrast, DNA methylation at certain genomic features such as enhancers is more dynamic (e.g., Argelaguet et al., 2019), and thus more variable across cells. Only the latter regions are of value for our goal of quantitating dissimilarity between cells. We call these the variably methylated regions (VMRs).

In the standard approach, one divides up (tiles) each chromosome into non-overlapping, equal-sized intervals, and quantitates the methylation of each tile. Such rigid placement of interval boundaries is unlikely to be optimal: for example, a VMR might be much smaller than a tile and the signal from its CpG sites will hence be drowned out by the larger number of uninformative CpG sites that are equal in all cells, when averaging over all the CpG sites in the tile.

Therefore, we propose the following approach (Fig. 2): Divide up the chromosome into many *overlapping* windows that start at regular multiples of a fixed, small, step size. Quantify the methylation of each cell in each window by averaging the cell’s methylation residuals over all CpGs in the window, as described above and depicted in Fig. 1B. Next, calculate for each window the variance of these values over all cells. Select, say, the top 2% windows with the highest variances and mark them as VMRs. Wherever thus marked windows overlap, merge them into one larger VMR. Then, calculate for each of these merged VMRs the methylation signal, as before, by averaging for each cell over the residuals of all contained CpG sites.

**Figure 2.**
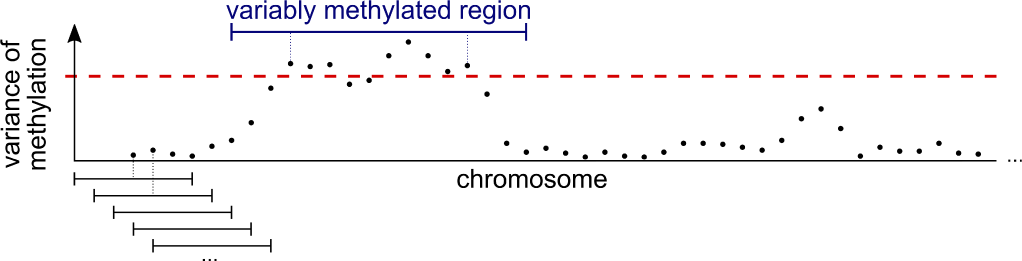
Finding variably methylated regions. The chromosomes are divided up into overlapping windows (first five shown at the bottom), and for each window, the cells’ methylation values are calculated as described and as depicted in Fig. 1B. Then, the variance of these values is calculated (each point represents one of the overlapping windows), a threshold (dashed line) is chosen such that a chosen quantile of windows have a variance exceeding the threshold. Windows with above-threshold variance are merged if they overlap, yielding the “variably methylated regions” (VMRs).

In this manner, we obtain a methylation matrix, with one row per cell and one column per VMR, that is, in a sense, richer in information and has better signal-to-noise ratio than the matrix obtained by the simple analysis sketched at the very beginning. As we demonstrate below, a PCA performed on such a matrix provided a distance metric for the cells that contains more information on biological detail than one from a simpler analysis.

### Application and benchmarks

To demonstrate the value of our proposed improvements, we benchmarked various combinations of analysis methods on five diverse single-cell methylome data sets, starting with a data set from our own research.

#### Correlating VMR methylation with gene expression

Our data set (Kremer et al., 2022) comprises the single-cell methylomes of 1566 cells isolated from mouse forebrains, as well as matched single-cell transcriptomes of the same cells. Among these cells are distinct cell types such as oligodendrocytes, oligodendrocyte precursor cells (OPCs) and endothelial cells, as well as cellular sub-states that are part of the continuous neural stem cell differentiation trajectory. To assess whether our VMR detection method captures genomic intervals that are biologically meaningful, we probed whether their methylation level correlates with the expression of nearby genes.

We first note that gene expression is more strongly correlated with the methylation of nearby VMRs than with the methylation of their promoters (Fig. 3A), indicating that VMR methylation is often a better predictor of gene expression than promoter methylation. Indeed, a gene-wise comparison revealed many genes whose expression is correlated with the nearest VMR, but not with promoter methylation. One such example gene, *Htra1*, is depicted in Fig. 3B. While the promoter of this gene is lowly methylated regardless of gene expression, a VMR located downstream of the promoter is lowly methylated in cells with high *Htra1* expression.

**Figure 3.**
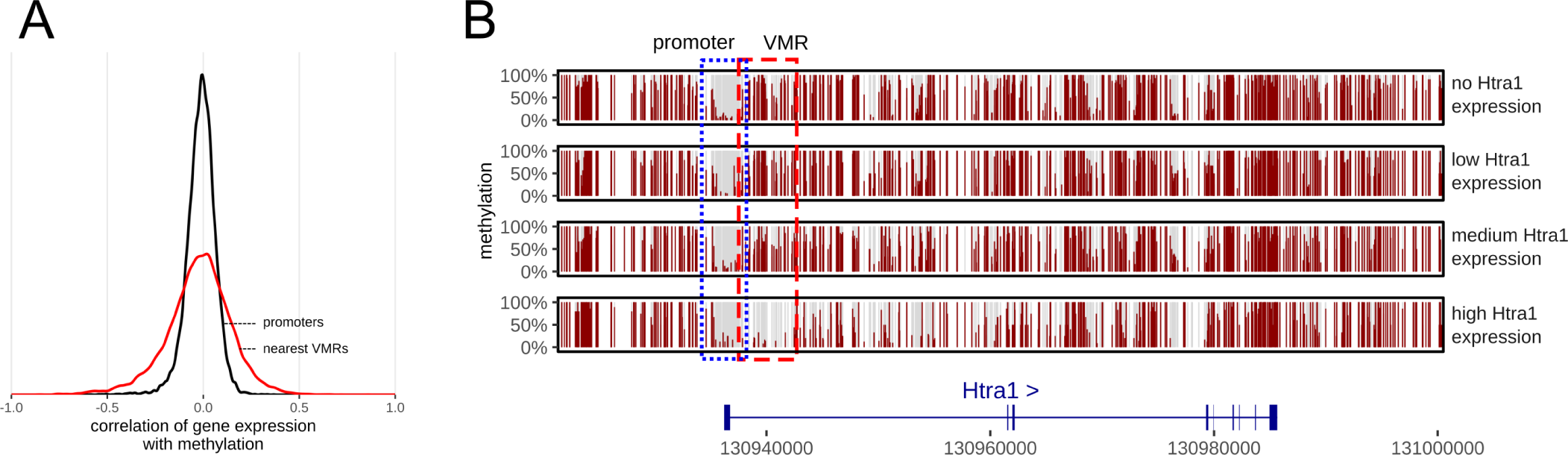
Correlation of DNA methylation and gene expression. **(A)** Distribution of Pearson correlations between gene expression and promoter methylation (black) and gene expression and methylation of the nearest VMR (red): for most promoters, correlation is weak. Promoters are defined as intervals ±2 kb around the TSS. **(B)** Mean methylation near the gene *Htra1*. Cells are assigned to 4 groups based on *Htra1* expression (group 0: cells with no *Htra1* expression, groups 1-3: cells that express *Htra1*, divided into three equally large groups with group 3 having the highest expression). Data from Kremer et al. (2022).

#### Improved identification of cell types

We next tested whether our methods improve the ability to distinguish cell types and cell states (Fig. 4). To this end, we obtained cell type/state labels based on the single-cell transcriptomes from Kremer et al. (2022) (Fig. 4A). We consider these cell labels as ground truth and tested whether we are able to distinguish the same groups of cells based on their methylomes. To do this, we subjected the methylomes to various combinations of analysis methods including our own proposed methods and others that are commonly used. Specifically, we selected four different sets of genomic intervals at which CpG methylation is to be quantified: either VMRs detected with our approach, 100 kb genomic tiles, promoter regions (i.e., transcription start site (TSS) ± 2000 bp), or candidate cis-regulatory elements (ENCODE cCREs, Moore et al., 2020). To quantify methylation at these features, we either simply averaged in these intervals, obtaining methylation percentages, or we calculated the shrunken mean of residuals, as described earlier. Finally, the resulting methylation matrices were subjected to iterative PCA (see Methods) and subsequent UMAP for visualization.

**Figure 4.**
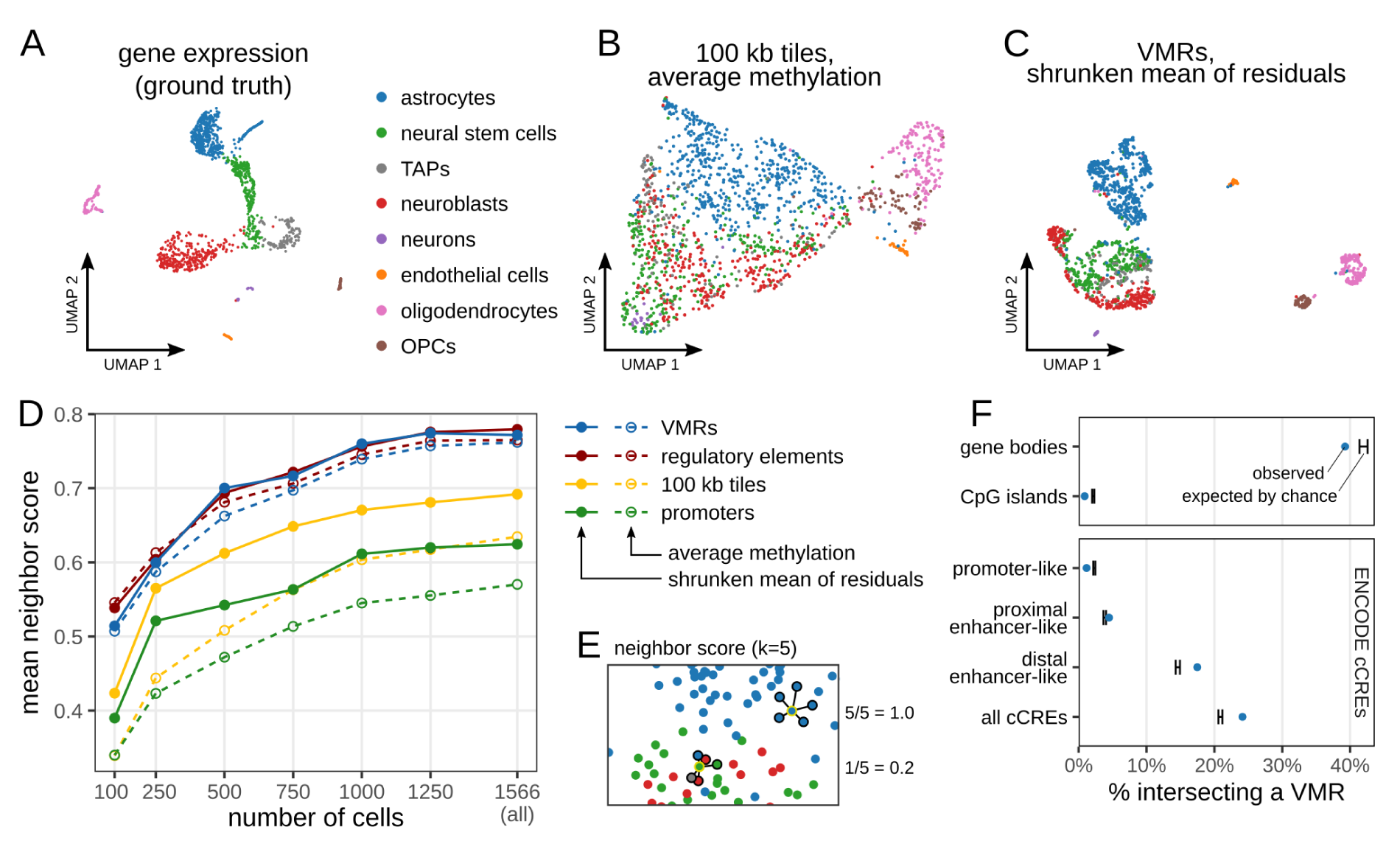
Benchmark of our methods on single-cell multi-omic data of cells of the murine forebrain. **(A)** Cell labels based on clustering of single-cell transcriptomes from Kremer et al. (2022). **(B, C)** Exemplary UMAPs obtained when analyzing the data set with conventional methods based on genome tiling (B) or our proposed methods (C). **(D, E)** Mean degree of cell type separation (neighbor score, E, computed in 15-dimensional PC space) obtained when analyzing single-cell methylomes with different combinations of methods. Either VMRs, ENCODE regulatory elements, 100 kb genomic tiles or promoter regions (TSS±2kb) were subjected to iterative PCA and UMAP. CpG methylation in these intervals was either quantified by averaging (dotted lines) or using the shrunken mean of the residuals as proposed in this work (solid lines). The full 1566-cell data set was sub-sampled to simulate smaller data sets (x axis). A higher neighbor score implies better separation of cell types inferred from single-cell transcriptomes of the same cells (A). **(F)** Proportion of VMRs (blue dots) that have at least 1 bp overlap with other genomic features. The black range illustrates the minimum and maximum overlap observed for 100 reshufflings of the VMRs to randomly chosen positions.

Visual inspection of the resulting UMAPs revealed that our proposed combination of methods results in more clearly separated cell types, compared to a UMAP obtained with default analysis methods (Fig. 4B,C). While all cells form a continuous point cloud when using default methods, our improvements led to a clear separation of oligodendrocytes, OPCs and endothelial cells. Furthermore, even cellular sub-states of cells in the continuous neural stem cell lineage were partially separated.

To quantify this performance gain in a more rigorous manner, we used a score that quantifies whether cells were placed, in 15-dimensional PC space, in a neighborhood comprising cells of the same cell type (“neighbor score”, see Fig. 4E, and Methods for details). The neighborhood relation in PCA space is relevant as it is used in many downstream analyses, e.g., for clustering. We ask how the neighbor score depends on the number of available cells. This is relevant as scBS protocols are costly and labor-intensive, and only few laboratories are currently able to obtain thousands of single cell methylomes. To simulate smaller data sets, we sub-sampled the 1566-cell data set into smaller data sets, analyzed them again with all combinations of analysis methods, and calculated the mean neighbor score for each (Fig. 4D). The results confirmed that quantifying methylation at VMRs leads to a cleaner separation of cell states than using promoter regions or 100 kb tiles. Using the shrunken mean of residuals as a measure of DNA methylation improved results further. This effect was most noticeable when quantifying promoters or genomic tiles, presumably because an individual promoter region or tile might span genomic regions with varying levels of DNA methylation as depicted in Fig. 1B, which our method accounts for. Although cell cluster separation generally becomes more difficult in those, performance gains were observed also in smaller data sets.

Overall, VMR quantification yielded results similar to those obtained when quantifying ENCODE regulatory regions, even though the number of detected VMRs (63 421) is considerably smaller than the number of ENCODE cCREs (339 815). When we repeated our analysis using only the 63 421 cCREs with the highest coverage the ability to distinguish cell types was diminished, suggesting that the average VMR is more informative for this task then the average cCRE (Fig. S3A,B). This demonstrates our *de novo* VMR detection approach’s ability to identify in scBS data a parsimonous set of relevant elements. The overlap between VMRs and cCREs is limited (Fig. 4F), indicating that VMR detection yields information that is complementary to other epigenetic marks. An further benefit of VMR detection is that this approach is available even in the absence of such annotations, for instance when studying species other than human or mouse. Lastly, using VMRs over regulatory regions resulted in decreased RAM requirements as well as a shorter runtime, even when accounting for the additional step of VMR detection (Fig. S3C,D).

We repeated this benchmark on an additional three published single-cell methylome data sets (Fig. S1). These include neuronal sub-types of the murine cortex (Luo et al., 2017, using cell type labels derived from CH-methylation in genomic tiles instead of CpG-methylation as ground truth), cells isolated from mouse embryos during the onset of gastrulation (Argelaguet et al., 2019, using RNA-derived cell clusters or alternatively embryonic stage as ground truth) and human colorectal cancer cells (Bian et al., 2018, using sampling region as ground truth). Again, we subjected each data set to all possible combinations of genomic feature selection and methylation quantification. We furthermore included three additional approaches to perform dimensionality reduction in our benchmarks, including PCA with two different pre-processing strategies, as well as MOFA+, a dimensionality reduction technique designed for multi-modal single-cell data that can also process methylation data (Argelaguet et al., 2020).

These extensive benchmarks confirmed that our proposed combination of methods, i.e. using the shrunken means of residuals of VMRs for dimensionality reduction, is able to distinguish diverse cellular properties such as cell type, colorectal cancer stage (normal tissue, primary tumour, metastasis), embryonic stage, and germ layer.

#### Robustness to parameter changes

Next, we assessed whether our proposed workflow requires fine-tuning of parameters (Fig. S2). To this end, we re-analyzed two data sets, as well as sub-samples of the data with different VMR detection parameters namely the width of the sliding window in bp, (set with option `--bandwidth` in our *MethSCAn* software, default 2000), the variance threshold above which windows are merged to VMRs (`--var-threshold`, default 0.02) and the step size of the sliding window (`--stepsize`, default 100bp). This parameter sweep showed that our workflow gives good results over a wide range of parameter values. For the CpG-methylation data of Luo et al. (2017), results are nearly independent of the parameters (Fig. S2B). In the more challenging data set of Kremer et al. (2022), cell types were less cleanly separated when very large bandwidths, very strict variance thresholds, or a very large step size was selected (Fig. S2A,C). However, very small bandwidths or very lenient thresholds resulted in a much higher number of VMRs and thus long computing times. Overall, our default parameter combination provided good results and fast compute times in both data sets.

#### Further applications

Lastly, we asked whether our methods are also suitable for the analysis of DNA methylation outside the default CpG context. To this end, we revisited the Luo et al. (2017) data set, but this time only considered CH-methylation (mCH). VMR detection with default options produced results that were qualitatively similar to those reported in Luo et al. (2017), suggesting that our methods are also suitable for this data type (Fig. S4A). Finally, as single-cell methylome data sets are expected to rapidly grow in size in the coming years, we furthermore performed a stress test on a large data set comprising 100 350 cells (Liu et al., 2021) (Fig. S4B).

### Finding differentially methylated regions (DMRs)

A common task in the analysis of bulk bisulfite-sequencing data is the detection of differentially methylated regions (DMRs) between conditions, tissues, or cell types (Hebestreit et al., 2013; Korthauer et al., 2018). As DNA methylation affects gene expression, DMRs can provide insights into the unique epigenetic and gene regulatory characteristics of cell types. However, to date no approach to detect DMRs in scBS data has been reported. To enable DMR detection in scBS data, we thus developed an approach that detects DMRs of variable size between two groups of cells, and controls the false discovery rate (Fig. 5):

**Figure 5.**
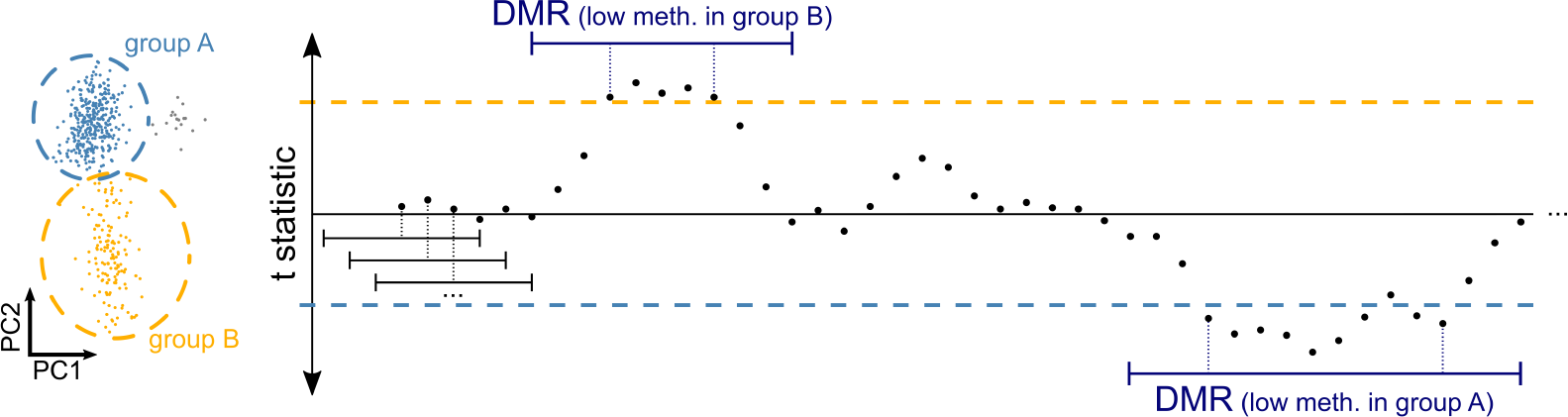
Identification of differentially methylated regions (DMRs). The chromosome is divided into windows overlapping by a small and fixed size. For each window, the cell’s methylation values are calculated as described and as depicted in Fig. 1B. Then the t statistic of these values is obtained as a measure of methylation difference between two groups of cells (each point represents one of the windows). Upper and lower thresholds (dashed lines) are chosen such that a defined quantile of windows have a t statistic above or below the threshold. Windows exceeding either threshold are merged if they overlap, yielding the differentially methylated regions (DMRs).

Similar to the previously described approach for VMR identification (Fig. 2), we divide each chromosome into overlapping windows shifted by a small and fixed size (step size) and quantify the methylation of each cell in each window. Next, instead of the variance, we obtain the t statistic as a measure of differential methylation between the two cell groups. We identify the windows with the most extreme t statistics, e.g. windows in the 2% upper and lower tails. We then merge any overlapping windows in the upper tail into larger DMRs, and do the same for windows in the lower tail, and then we re-calculate the t statistic for each larger DMR.

To assess statistical significance of DMRs, we repeat the same procedure on permutations of the scBS data, i.e. the same data set with randomly shuffled cell labels. The DMR t statistics obtained from permuted data are then used to estimate the false-discovery rate (FDR), yielding an adjusted p-value for each DMR.

While the primary purpose of VMRs is to provide better input for PCA and distance calculations, DMR detection facilitates the discovery of epigenetic differences between conditions or cell types, as we demonstrate next.

### Detecting DMRs between oligodendrocytes and neural stem cells

We used the single-cell multi-omics data set (Kremer et al., 2022) to evaluate our DMR detection approach. We first selected two cell populations from the healthy murine ventricular-subventricular zone: neural stem cells (NSCs, 130 cells) and oligodendrocytes (58 cells). Then we used the method just described to identify DMRs between these two cell types (Fig. 6A). Repeating this after permuting the cell-type labels, in order to obtain a null distribution of the t statistics, yielded DMRs with much weaker methylation differences. Consequently, we could assign to many of the DMRs detected in the unpermuted data a low adjusted p-value (colors in Fig. 6A).

**Figure 6.**
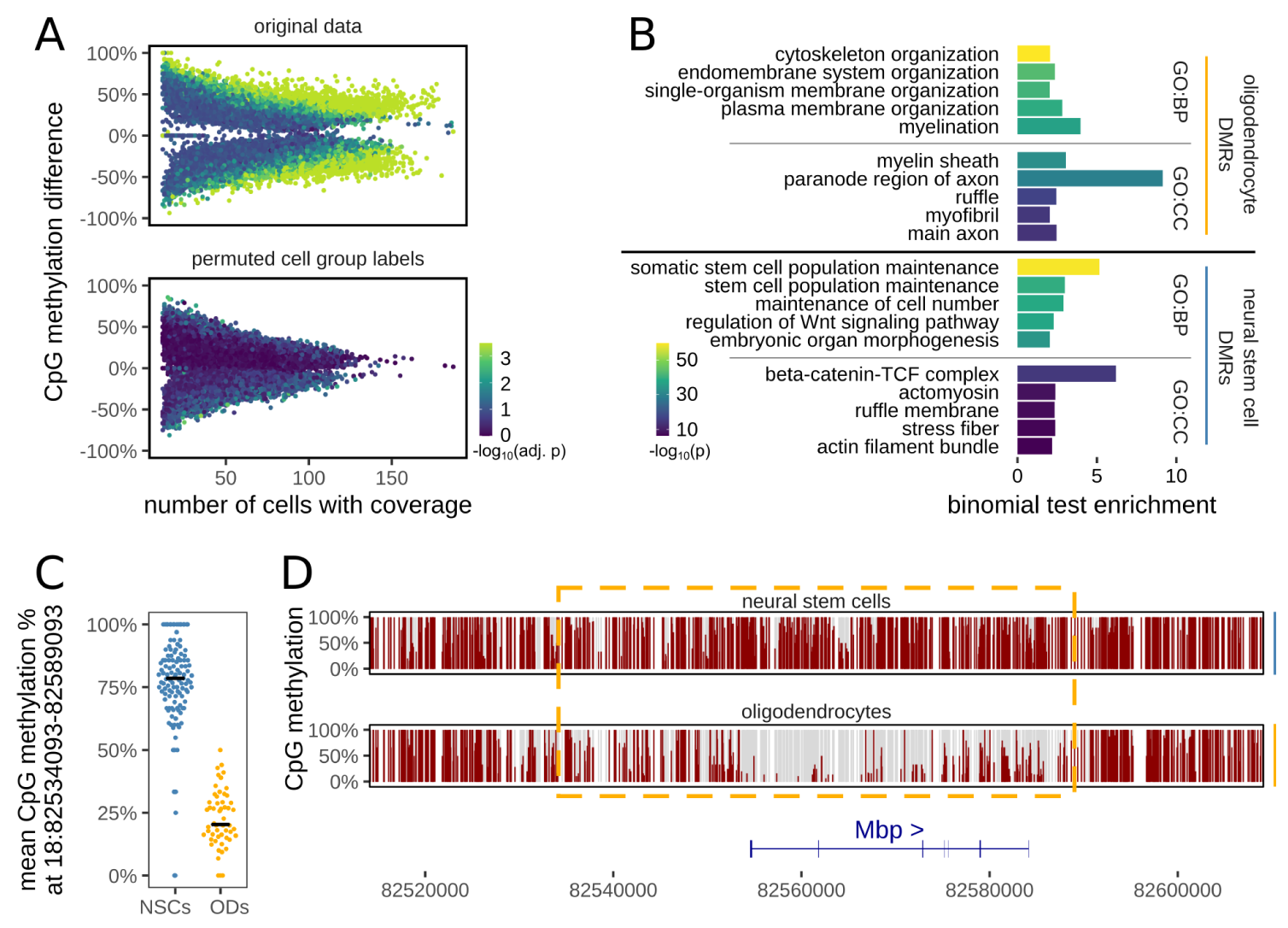
Detection of differentially methylated regions between oligodendrocytes and neural stem cells. **(A)** DMRs detected between 58 oligodendrocytes (ODs) and 130 neural stem cells (NSCs) from Kremer et al. (2022) (top) and DMRs detected in the same data with randomly permuted cell labels (bottom, used to estimate FDR and determine adjusted p-values). **(B)** Enrichment of GO terms associated with DMRs lowly methylated in ODs (top) or NSCs (bottom). Depicted are the top 5 GO terms of the “Biological Process” (GO:BP) and “Cellular Component” (GO:CC) GO category, and their binomial test p-value and enrichment, as reported by GREAT (McLean et al., 2010). **(C)** Mean methylation of NSCs and ODs at an exemplary DMR. Each point corresponds to a cell, black lines denote the median. **(D)** Detailed view of the DMR (yellow dashed rectangle) from C in pseudo-bulk samples consisting of NSCs or ODs. Vertical bars represent CpG sites.

Gene ontology (GO) enrichment with GREAT (McLean et al., 2010) revealed that DMRs lowly methylated in oligodendrocytes are located near genes involved in myelination, the main function of oligoden-drocytes. Similarly, DMRs specifically demethylated in NSCs occur near genes involved in stem cell population maintenance. This demonstrates that our DMR detection approach is able to identify biologically meaningful DMRs, even in scBS data sets of modest cell number.

Fig. 6C-D depicts an exemplary DMR, located at the gene body of myelin-basic protein (*Mbp*), the major component of myelin that is essential for myelination of neuronal axons (Boggs, 2006). Our results suggest that oligodendrocyte-specific gene expression of *Mbp* is supported by low methylation at the detected DMR.

### The *MethSCAn* software toolkit

We have implemented the approach just described in a Python package with a command line interface, *MethSCAn* (ANalysis of Single-Cell METHylation data), which also offers a number of other functionalities for the analysis of scBS data (Fig. 7). The starting point of such an analysis are methylation files generated by tools such as Bismark (Krueger and Andrews, 2011), methylpy (Schultz et al., 2015) or BISCUIT (Zhou et al., 2021). For each cell, these tools produce a text file that lists the methylation status of all CpG sites. Since it is inconvenient to work with hundreds or thousands of text files, *MethSCAn* provides the prepare command which parses these methylation files and stores their content in a compressed format that enables efficient access to all CpG sites of the genome. `methscan prepare` is flexible and accepts all tabular input formats including Bismark .cov-files, methylpy .allc-files, or custom user-defined formats.

**Figure 7.**
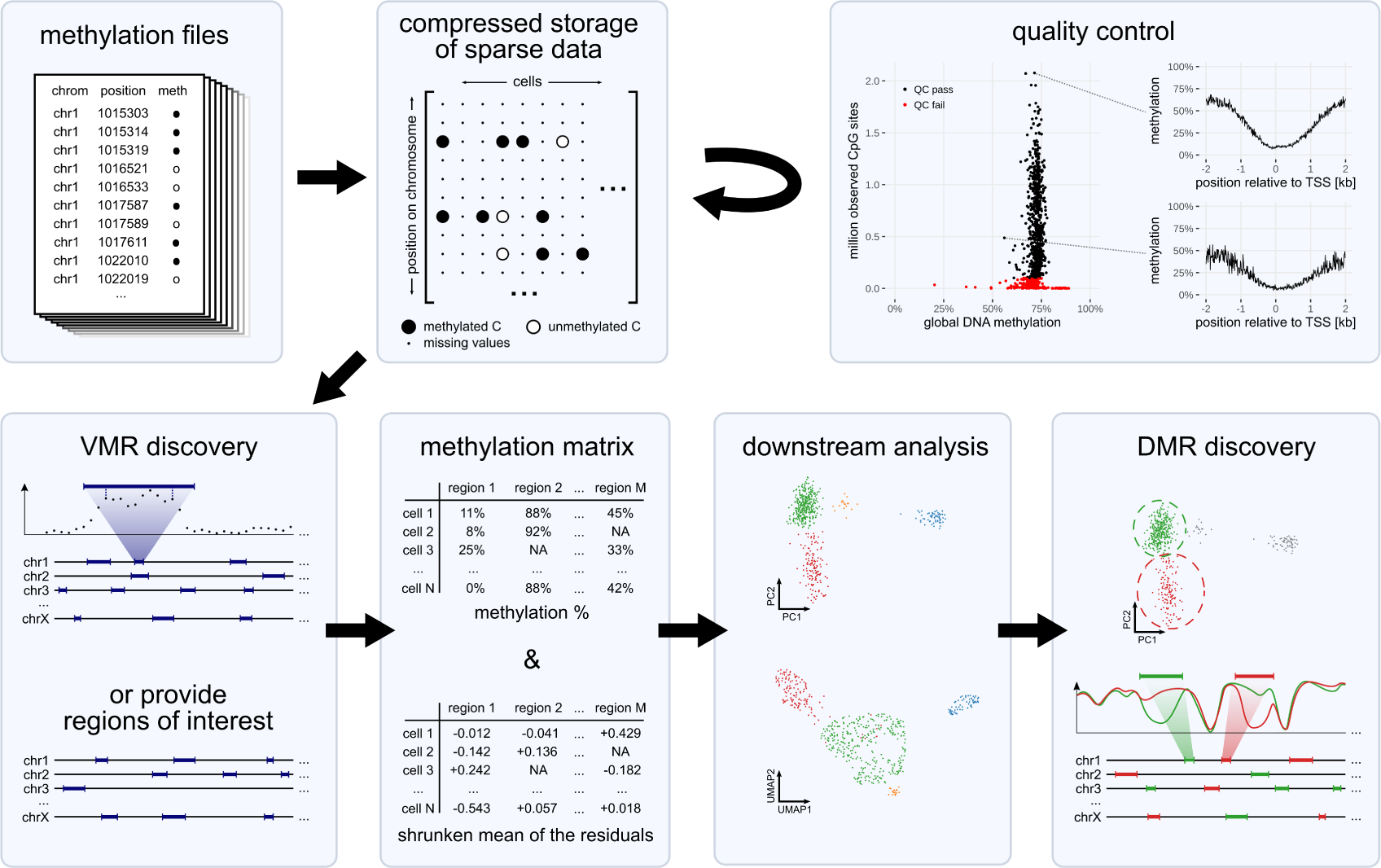
Overview of the functionalities implemented in the *MethSCAn* package. `methscan prepare` parses methylation files produced by common bisulfite sequencing mappers and stores their contents in a compressed format optimized for efficient access to genomic intervals. *MethSCAn* also produces cell-wise summary statistics and quality plots (here: average methylation around the transcription start site) that are used to detect low-quality cells. These cells can be discarded with `methscan filter`. To obtain a methylation matrix, similar to the count matrices used in scRNA-seq, the user must first decide in which genomic intervals methylation should be quantified. The user may either provide genomic regions of *a priori* interest, or they may choose to discover VMRs (variably methylated regions) in the data with `methscan scan`. The resulting methylation matrix can then be used for downstream analysis such as cell clustering and dimensionality reduction. Identified groups of cells (e.g. clusters) can then be scanned for DMRs with `methscan diff`.

In brief, we store methylation of each chromosome in a matrix where each column represents a cell and each row represents a base pair. Methylated and unmethylated sites are coded as +1 and − 1, respectively. Since the vast majority of values in this matrix are missing due to the sparsity of scBS data (and because rows for base-pairs not corresponding to a CpG site contain no data), we encode missing values as zero and then store the data in Compressed Sparse Row (CSR) format. This format does not explicitly store zeroes (here: missing values) and is optimized for row-wise access, which results in significant compression and allows fast access to the methylation status of genomic intervals.

methscan prepare also computes a number of summary statistics for each cell, including the mean genome-wide methylation level and the number of observed CpG sites, i.e. the number of CpG sites that have sequencing coverage. These summary statistics can be used to detect cells with poor quality. The quality of each single cell methylome can furthermore be inspected with `methscan profile`, which computes the average methylation profile of a set of user-defined genomic regions such as transcription start sites (TSS) at single-cell resolution. The TSS profile is a useful quality control plot since methylation shows a characteristic dip roughly ±1 kb around the transcription start site in mammalian genomes. Cells that do not show this pattern, or cells with few observed CpG sites, may then be discarded with `methscan filter`.

After quality control, the user has access to the genome-wide VMR detection approach described earlier, using the `methscan scan` command. This produces a BED file that lists the genome coordinates of VMRs. To finally obtain a methylation matrix analogous to a scRNA-seq count matrix, this BED file can be used as input for `methscan matrix`, which quantifies methylation at genomic intervals in all cells. The command produces both, the simple percentage of methylated CpG sites, as well as our proposed methylation measure, the shrunken mean of the residuals, which is more robust to variation in sequencing coverage or stochastic differences in read position between cells. We note that both `matrix` and `profile` accept any valid BED file as input, which means that the user can quantify and visualize methylation at any set of genomic features of interest, such as promoters, enhancers or transcription factor binding sites, obtaining one profile plot per cell. The methylation matrix produced by the `matrix` function can then serve as input for established methods used for the exploration of single-cell data, such as dimensionality reduction and cell clustering.

Finally, after annotation and exploration of the data set, the user may specify two groups of cells for DMR detection with `methscan diff`. This command produces a BED file listing the genome coordinates, adjusted p-values, as well as several other metrics associated with DMRs. These DMRs may then be associated with nearby genes, or used for GO enrichment with tools such as GREAT (McLean et al., 2010) to enable functional interpretation.

## Discussion and Conclusion

Here, we have proposed an improved strategy to preprocess single-cell bisulfite sequencing data. Based on the observation that incomplete read coverage of a genomic interval can lead to inaccurate methylation estimates, we suggest a scoring scheme that is aware of a read’s local context rather than just treating all reads in an interval alike.

Furthermore, we show a way to pinpoint minimal regions of high variability across cells, which we call variably methylated regions (VMRs). Unlike other tools for scBS data analysis (Kapourani and Sanguinetti, 2019; Kapourani et al., 2021; Danese et al., 2021), which rely on the user to manually specify which genomic intervals should be quantified, *MethSCAn* implements an approach to discover these intervals in the data itself. This not only reduces noise and allows to focus only on the features that are important for the given dataset, but also provides useful input for genomics-style analyses. Depending on the research question at hand, individual VMRs may be related to nearby genomic features such as gene bodies or known regulatory elements, or subjected to gene ontology and motif enrichment.

To furthermore aid interpretation of scBS data, we developed and implemented an algorithm for genome-wide detection of DMRs in single-cell methylomes. FDR estimation via permutation allows us to report statistical significance of each DMR. In a similar manner as VMRs, pinpointing the regions that differ between groups of cells and hence have a putative regulatory role aids biological interpretability. For instance, applying our approach to NSCs and oligodendrocytes demonstrated that the obtained DMRs locate near meaningful loci associated with cell-type specific functions.

We also presented and published an open source software tool, called *MethSCAn*, that provides an easy-to-use implementation of the described algorithms. It can start directly from the output of methylation callers such as Bismark (Krueger and Andrews, 2011), Biscuit (Zhou et al., 2021) and methylpy (Schultz et al., 2015) and produce a cell×region matrix. We suggest iterative PCA as an approach to map the count matrix to a reduced-dimensional space overcoming the abundance of missing values. Alternatively, one may use for this purpose established tools based on matrix factorisation such as MOFA+, (Argelaguet et al., 2020, included in our benchmarks) or LIGER (Welch et al., 2019). Once a low dimensional embedding is obtained, one can switch to well established methods from scRNA-seq analysis, including visualization tools such as t-SNE and UMAP, Leiden/Louvain clustering, pseudotime trajectory analyses, etc., e.g., by using either overall toolkits like Seurat (Hao et al., 2023), ScanPy/EpiScanPy (Wolf et al., 2018; Danese et al., 2021) or any of the many available tools for specific tasks. By offering these functions, *MethSCAn* bridges a gap in the chain of existing tools that so far hindered practitioners in their data analysis. Our implementation can handle datasets of various sizes up to a hundred thousand cells (see Fig. S4B).

By re-analyzing four published data sets, we showed that these improvements to data preprocessing help to increase signal and decrease noise, resulting in a more informative intermediate-dimensional representation of the data. As examples of practical benefits, we demonstrate that our preprocessing allows for better distinction of cell sub-types, especially for challenging data sets comprising cellular sub-states and lineage transitions or for data sets with small cell number.

In conclusion, we have presented powerful improvements to scBS data preprocessing and a software tool that implements these.

## Data Availability

Single-cell multi-omic data of cells of the murine fore-brain (Kremer et al., 2022) is available at the NCBI Gene Expression Omnibus (GEO) under the accession GSE210806.

## Code Availability

*MethSCAn* is free and open source. The package is available on the Python Package Index (PyPI) and can be installed with the Python package installer pip. The source code is hosted at https://github.com/anders-biostat/MethSCAn, where we also provide detailed documentation including a tutorial that demonstrates a complete *MethSCAn* analysis on an example data set.

## Methods

### Raw data

Let us write *x*_*ij*_ for the methylation status of CpG *i* in cell *j*. The index *i* runs over all CpG positions present in the genome, the index *j* over all cells in the assay. We write *x*_*ij*_ = 0 if position *i* was found to be unmethylated in cell *j* by bisulfite sequencing, *x*_*ij*_ = 1 if it was methylated, and *x*_*ij*_ = NA if position *i* was not covered by reads from cell *j* and the methylation status is therefore not available (“NA”).

These values can be readily obtained from single-cell bisulfite sequencing data using tools like Bismark.

If multiple reads from the same cell cover a position, these will typically be PCR duplicates of each other and hence agree. Of course, the two alleles of a CpG may rarely differ in their methylation status. While it is in principle possible that one obtains discordant reads stemming from the same position on both the paternal and the maternal chromosomes of the same cell, this is so unlikely that we can ignore the case. Hence, whenever the methylation caller reports multiple reads covering the same position in the same cell, we set *x*_*ij*_ to 0 or 1 whenever all reads agree. When there is disagreement, we put *x*_*ij*_ = NA by default, or optionally follow the majority of reads whenever possible.

For later use: We define *C* as the set of all cells in the assay (i.e., *C* is the index set for the cell indices *j*). Moreover, we define *C*_*i*_ ⊂ *C* as the set of all those cells *j* that have reads covering position *i*,

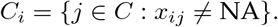

Conversely, we define *G*_*j*_ as the set of all the CpG positions *i* covered by reads from cell *j*

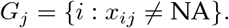

### Data storage

The function `methscan prepare` reads a set of methylation files (e.g. produced by Bismark) and produces a file that stores the matrix *x* in a space-efficient format, as follows: *x* is represented as a SciPy sparse matrix (Virtanen et al., 2020), encoding the actual values 0, 1, and NA as -1, 1, and 0, respectively. Coding NA (the most common value) as 0 leverages SciPy’s sparse matrix storage. In all that follows here, any mention of *x* will, however, always mean the encoding as *x*_*ij*_ ∈ {0, 1, NA}.

### Smoothing

For each CpG position *i*, we write

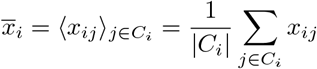

for the average methylation at position *i*, where⟨·⟩ denotes averaging, and the average runs over all the cells *j* ∈*C*_*i*_, i.e. over those cells for which methylation data is available for position *i*.

We then run a kernel smoother over these per-position averages to obtain the smoothed averages 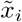. Specifically, we use

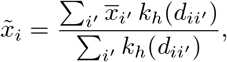

i.e., 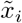 is the weighted average over the per-position averages 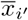, taken over the CpG sites *i*^′^ in the neigh-borhood of *i*, and weighted using a smoothing kernel *k*_*h*_ with bandwidth *h*. Here,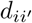 is the distance between CpG positions *i* and *i*^′^, measured in base pairs, *h* is the smoothing bandwidth in base pairs (by default, *h* = 1000), and *k*_*h*_ is the tricube kernel,

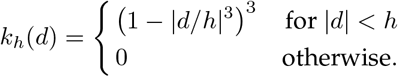

### Methylation for an interval

Next, we discuss averaging methylation over a range of CpG sites.

Given an interval *I* on the chromosome, we wish to quantify the average methylation *m*_*Ij*_ of the CpG sites within the interval for cell *j*. If we interpret *I* as the set of CpG positions *i* in the interval, we may write

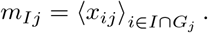

Here, the average runs over all those sites *i* that lie within the interval *I* and are covered by reads from cell *j*.

If we wish to compare cells, it can be helpful to center this quantity by subtracting its average:

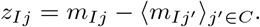

As an alternative, we suggest to consider the residuals of the individual CpG methylation values *x*_*ij*_ from the smoothed average 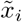,

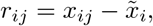

and averaging over these, obtaining

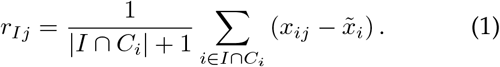

This is a shrunken average, with denominator *n* + 1. This extra pseudocount has the effect of shrinking the value towards the “neutral” value 0, with the shrinkage becoming stronger if the data is “weak” because the number | *I* ∩*C*_*i*_ | of positions covered by reads from cell *j* is low. In the extreme case of none of the reads from cell *j* covering *I*, the sum becomes 0 and the denominator 1, i.e. 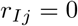 in this case.

### Finding variably methylated regions

For any interval *I*, we denote by *v*_*I*_ the variance of its residual averages 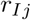:

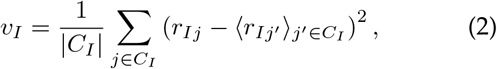

where the average runs only over the set *C*_*I*_ = ⋃_*i*_ ∈_*I*_ *C*_*i*_ of those cells *j* which have reads covering interval I.

To find VMRs, we define intervals *I*_1_, *I*_2_, …, all of the same width, and with step-wise increasing starts, then calculate *v*_1_, *v*_2_, … for these intervals. We then mark the intervals with the 2% highest variances. We take the union of all these intervals, split the union into connected components, and call each component a VMR. Putting that last step in other words: We take all the intervals with variance in the top 2-percentile, fuse intervals that overlap and call the regions thus obtained the VMRs.

### Finding differentially methylated regions

When searching for DMRs, we compare two groups of cells, whose index sets we denote with *C*_A_ and *C*_B_. For a given interval *I*, we calculate the mean each of the mean shrunken residuals *r*_*ij*_ (see Eq. (1)) over the cells *j* in each of the two groups,

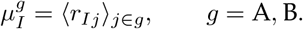

We also calculate a variance as in Eq.(2):

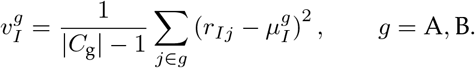

From this, we calculate Welch’s t statistic as usual:

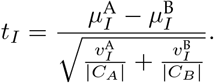

In order to find candidate DMRs, we again define overlapping and stepwise shifted intervals *I*_1_, *I*_2_, … as for the VMRs and calulate t statistics *t*_1_, *t*_2_, … for these. As before, we take the top 2-percentile of these values, fuse intervals that overlap and call the regions thus obtained candidate DMRs. We repeat the procedure for the bottom 2-percentile to get the candidate DMRs for the other sign.

Next, we need to check these candidate DMRs for statistical significance. We first remind the readers here that, as this is a within-sample analysis, and cells, not samples, are the statistical unit. Therefore, a call as significant implies that the same DMR is likely to be called again if we repeated the analysis with another set of cells taken from the same biological sample, not that it would generalize to further samples. This fact, although often overlooked, is common to all within-sample analyses in the single-cell fields, e.g. also to the differential expression tests performed in scRNA-seq analyses to find marker genes that differentiate clusters.

It may seem that we could use the standard procedure for the Welch t-test here, i.e. use the Welch-Satterthwaite formula to get an approximate degree of freedom and then calculate the tail probability of the corresponding t distribution. However, this is unlikely to hold for two reasons: First, the Welch-Satterthwaite degrees of freedom are only based on the number of cells per group and do not account for the fact that the read coverage might vary from cell to cell. Second, the fusing of the DMRs obtained in the scanning step to obtain fused candidate DMRs would invalidate subsequent p-value based adjustment for multiple testing.

Therefore, we have instead implemented a permutation procedure, which works as follows: We randomly shuffle the assignment of the cells in *C*_A_ ∪*C*_A_ to either of the two groups and then rerun the whole procedure, i.e., the scanning step, the DMR fusing and the calculation of t values from the (potentially fused) candidate DMRs. This needs to be done for a sufficiently large number of permutations. Running through the whole genome for each permutation would be too computationally expensive. Instead, we go through the genome only once, but reshuffle the group labels every 2 Mbp.

All the t values obtained from this permutation procedure are taken together to get an empirical null distribution. Then, we can use this null distribution to control false discovery rate (FDR) by applying the Benjamini-Hochberg procedure in its p-value-free form: Let us write *T* for the set of all t values obtained from the “real” assignment of cells to group labels and *T*_0_ from the set of all t values obtained from the shuffled assignments, i.e., the empirical null distribution. To adjust a specific t value *t*_*i*_ ∈ *T*, we calculate

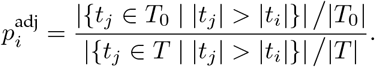

In words: we calculate which fraction of the null t values is greater than t by absolute value, and which fraction of the real t values is. The ratio gives us the false discovery rate we should expect if we used the t value as threshold.

### Calculating cell to cell distances

Given a set, 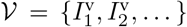, of intervals corresponding to VMRs, we get a relative methylation fraction *r*_*ij*_ for each VMR 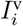 and each cell *j* from Eq. (1). The matrix thus obtained can then be centered and used as input for a PCA. If we calculate the top *R* principal components, we thus obtain for each cell *j* an *R*-dimensional principal component vector 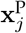. For any two cells *j, j*^′^, we use the Euclidean distance 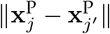 as the measure of dissimilarity of the two cells. Thus, the matrix of PC scores can be used as input to dimension reduction methods like t-SNE or UMAP, and to clustering methods like the Louvain or Leiden algorithm, which require such a matrix as input to the approximate nearest neighbor (ANN) finding algorithm that is their first step.

### PCA with iterative imputation

Whenever a region is not covered by any read in a cell, the corresponding data entry in the input data matrix for PCA will be missing. The standard approach to calculate PCA, commonly done using the IRLBA algorithm (Baglama and Reichel, 2005), is not suited to deal with missing data. We circumvent this issue by simply using the PCA itself to impute the missing value in an approach that we call “iterative PCA”).

Let us write *A* for the matrix to which the PCA is to be applied, with the features (here: regions) represented by the matrix rows. The matrix has already been centered, i.e., Σ_*i*_ *a*_*ij*_ = 0. To establish notation, we remind the reader that performing a PCA on A means finding the singular value decomposition (SVD), *A* = *UDR*^⊤^, with *D* diagonal and *U* and *R* orthonormal. The PCA scores are contained in *X* = *UD*, the loadings in *R*. To reconstruct the input data *A* from the PCA representation, one may use *A* = *XR*^⊤^, i.e., *a*_*ij*_ = Σ_*r*_ *x*_*ir*_*r*_*jr*_, where the equation is exact if *r* runs over all principal components and approximate if it is truncated to the leading ones.

Our iterative imputation strategy is now simply the following: we first replace all missing values in the row-centered input matrix *A* with zeroes and perform the (truncated) PCA. Then, we use the PCA predictions for the missing values, i.e., the *a*_*ij*_ = Σ_*r*_ *x*_*ir*_*r*_*jr*_, as refined stand-ins for the missing values in *A* and run PCA once more. This can now be iterated until convergence.

We note that similar approaches have also been used elsewhere (Josse and Husson, 2012).

### Analysis of scBS datasets for benchmarks

To analyze scBS data from Kremer et al. (2022), single-cell CpG methylation reports from all conditions were first stored with `methscan prepare` and then smoothed with `methscan smooth` using the default bandwidth of 1000 bp. This data was then analysed multiple times with different combinations of analysis methods, namely four ways to divide the genome into intervals, two ways to quantify methylation in these intervals, and four approaches for dimensionality reductions. The following four sets of genomic intervals were used: VMRs, obtained with `methscan scan` using the current default options (bandwidth = 2000, stepsize = 10, variance threshold = 0.02, minimum cell requirement = 6); adjacent tiles of 100 kb width; promoter regions as defined by the ±2 kb domain around the TSS of coding genes; and candidate cis-regulatory regions annotated by the ENCODE consortium (Moore et al., 2020). Methylation was quantified either by averaging binary methylation values, or by calculating the shrunken mean of the residuals as described earlier.

We used four different approaches for dimensionality reduction. Three of them involve imputation of missing values followed by PCA: The first approach, iterative PCA, was described earlier. Second, “PCA on highquality features” imputes missing values with the mean methylation level of a the given interval, while only retaining high-quality features selected as in Luo et al. (2017): We selected tiles spanning ≥ 20 CpG sites and with sequencing coverage in at least 70% of cells. We then imputed missing values with the mean of each tile, centered the values, and performed PCA. The third approach, “mean-imputed PCA” is identical to approach two, but without the quality-filtering step. Lastly, we used MOFA+ with default parameters instead of PCA, which does not require imputation of missing values. In all four cases we reduced the dimensionality of the input data to 15 PCs or MOFA factors. In some cases MOFA+ returned a smaller number of factors, since some of the requested 15 had zero variance. For visualization, these 15 PCs or factors were subjected to UMAP with parameters min dist = 0.2, init = “spca”. To flexibly adapt to data sets of different sizes, we set n neighbors 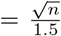 (rounded to the nearest integer), where *n* is the total cell number. The same analysis was repeated for three additional scBS data sets (Luo et al., 2017; Bian et al., 2018; Argelaguet et al., 2019), and for smaller data sets generated by randomly sub-sampling cells separately from these data sets.

VMRs that intersect protein-coding gene bodies, CpG islands (from the UCSC genome browser) or cCREs were quantified by subtracting VMRs with at least one bp of overlap using bedtools subtract -A (Quinlan and Hall, 2010) and counting the remaining VMRs.

To test our DMR detection approach, we selected oligodendrocytes and NSCs from healthy wild type mice of Kremer et al. (2022) and ran methscan diff with default parameters. For GO enrichment analysis, DMRs with adjusted *p <* 0.01 were uploaded to GREAT (McLean et al., 2010) with the association rule “basal plus extension, 0 bp upstream, 20 kb down-stream, 1 Mbp max extension, curated regulatory domains included”.

### Mean neighbor score

To assess performance of our methods, we employed a score that quantifies how well cell types (or cell states) are separated in 15-dimensional PCA space. For data from Luo et al. (2017), we used cell type labels that the authors manually curated based on CH-methylation. For the multi-omic data set, we repeated the single-cell transcriptomics analysis described in Kremer et al. (2022) with two adjustments: We did not filter off-target cells such as endothelial cells, and we annotated cell types using Leiden clustering with the Seurat (Stuart et al., 2019) function `FindClusters(resolution = 0.1)`. The score, based on the Γ-score (Kireeva et al., 2014), varies between 0 and 1, where higher scores reflect better separation of cell types. For each cell *j*, we count how many of its *k* nearest neighbors have been assigned to the same cell type as cell *j*. We denote this count 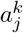 where we have chosen 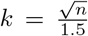, rounded. The overall score used to evaluate a given combination of methods is then simply the mean of all cell-wise scores.

### Correlation of DNA methylation and gene expression

To assess correlation between gene expression and the methylation status of VMRs or promoters, we first detected VMRs with `methscan scan --bandwidth 1000 --var-threshold 0.05`. We then quantified DNA methylation at VMRs and promoters with `methscan matrix`. We defined promoters as ± 2000 bp regions centered on a genes transcription start site (TSS). When multiple TSSs were annotated, we chose the TSS of the “principal” isoform (Rodriguez et al., 2013). Log-normalized expression values reported by Kremer et al. (2022) were then correlated with methylation of the gene’s promoter, or with methylation of the VMR closest to the gene body. When multiple VMRs intersected the gene body, we chose the VMR with the highest methylation variance. As a measure of methylation we used the shrunken mean of the residuals. We omitted lowly expressed genes (with scRNA-seq counts in *<* 5% of cells) and promoters and VMRs with low scBS coverage (in *<* 10 cells).

## Acknowledgments

SA and LPMK acknowledge funding by the Klaus Tschira Foundation (project 00.022.2019). LPMK, MMB, LK, SC, and AMV acknowledge funding from the European Commission via ERC grant 771376. We thank Alexey Uvarovskii for code refactoring and helpful suggestions on the source code.

## Supplementary Figures

**Figure S1:**
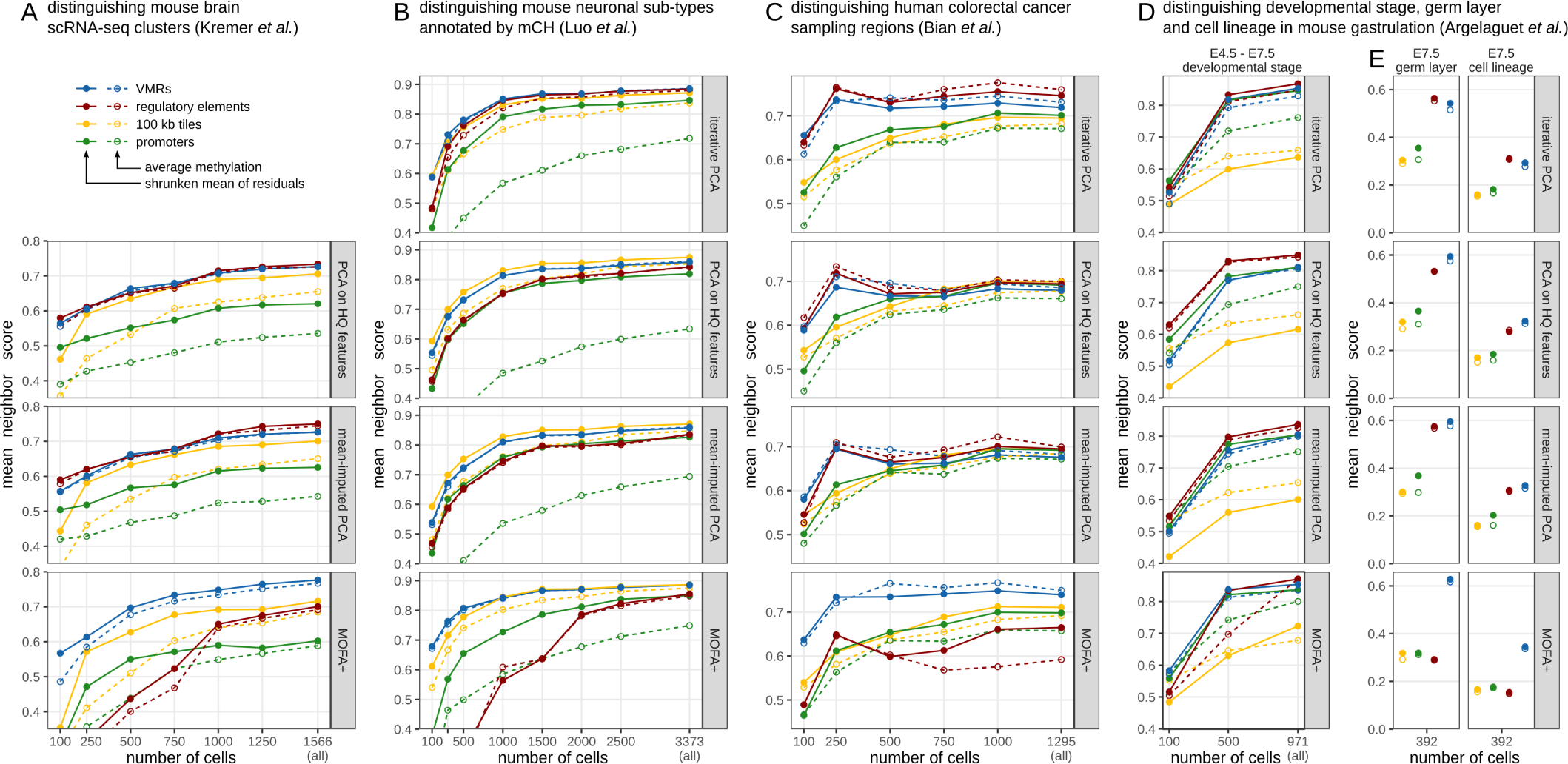
Benchmark of several combinations of analysis methods on four single-cell methylation data sets. **(A-E)** Benchmarking the ability to separate groups of cells based on CpG methylation data, using different analysis approaches. Methylation matrices were obtained either by averaging CpG methylation in genomic intervals (dotted lines, hollow points) or using the shrunken mean of residuals (solid lines and points). Each analysis was performed for the full data sets and sub-samples of each. Four sets of genomic intervals (VMRs, ENCODE regulatory elements, 100 kb tiles or promoters) were quantified and separately subjected to four dimensionality reduction techniques (see Methods for details): iterative PCA as proposed by us, mean-imputed PCA on intervals with high sequencing coverage (Luo et al., 2017), mean-imputed PCA on all intervals, and MOFA+ (Argelaguet et al., 2020) on all intervals. For each combination of methods, we quantified the ability to use distance in 15-dimensional PCA space to separate ground-truth cell labels reported by the authors (neighbor score). **(A)** Separation of neural cell types/states annotated based on scRNA-seq of the same cells (Kremer et al., 2022). **(B)** Separation of neuron sub-types, annotated by averaging CH-methylation (mCH) in 100 kb genomic tiles (Luo et al., 2017). **(C)** Separation of human colorectal cancer sampling regions (primary tumor, normal adjacent tissue, lymph node metastasis, liver metastasis before and after treatment, omental metastasis) (Bian et al., 2018). **(D-E)** Separation of three properties of mouse embryo cells: The developmental stage (E4.5, E5.5, E6.5, E7.5), the germ layer or extra-embryonic tissue of origin, or the cell lineage. Germ layer and lineage (E, only E7.5-cells) were annotated by Argelaguet et al. (2019) based on scRNA-seq of the same cells.

**Figure S2:**
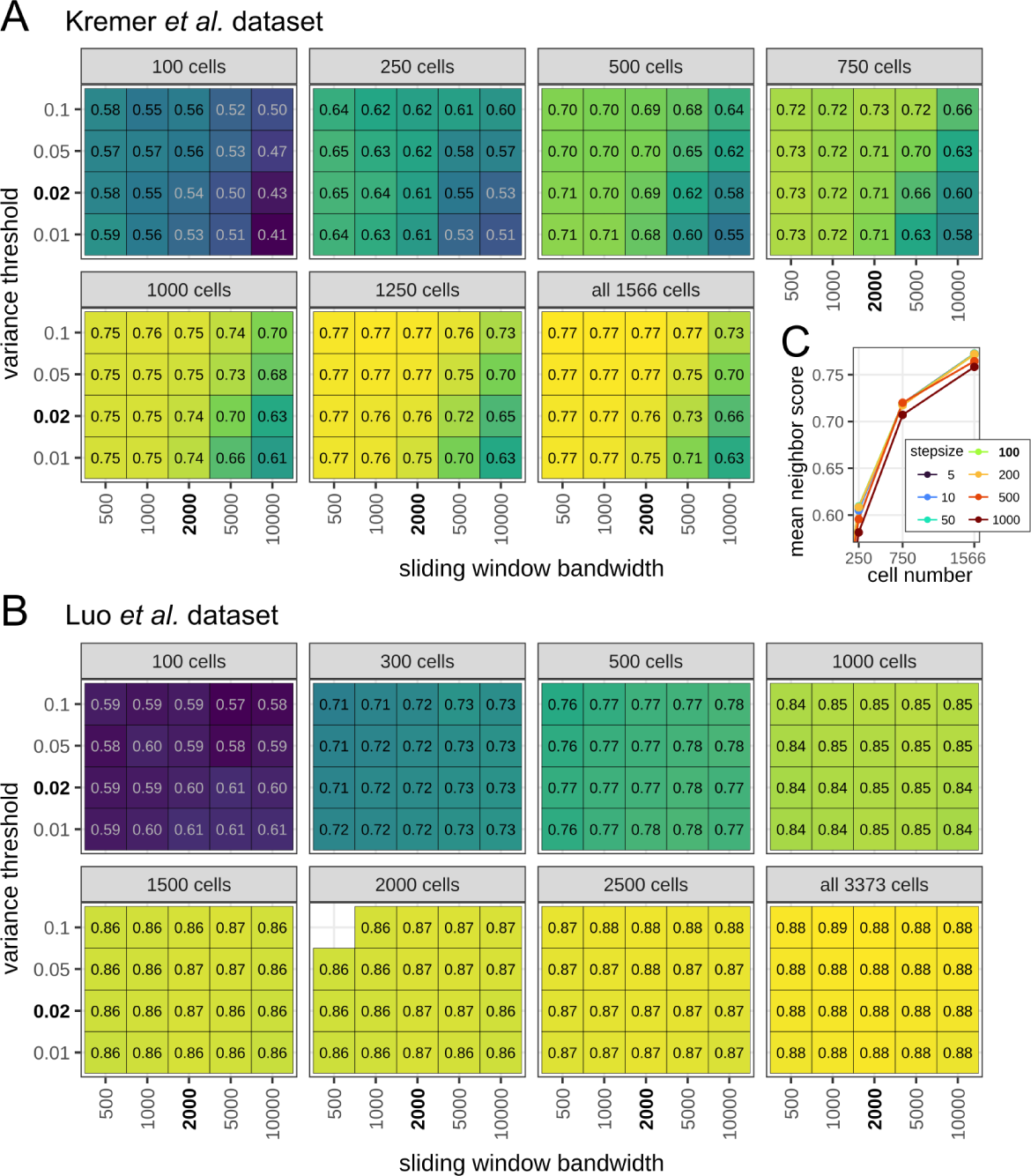
Effect of VMR detection parameters on separation of cell types. **(A-B)** Mean neighbor scores obtained after analyzing single-cell methylomes from Kremer et al. (2022) (**A**) or Luo et al. (2017) with our proposed methods. VMRs were detected with `methscan scan` using various sliding window bandwidths, variance thresholds, and on sub-samples of the full data sets. **(C)** Mean neighbor scores obtained after analyzing the Kremer et al. (2022) data set with various sliding window step sizes.

**Figure S3:**
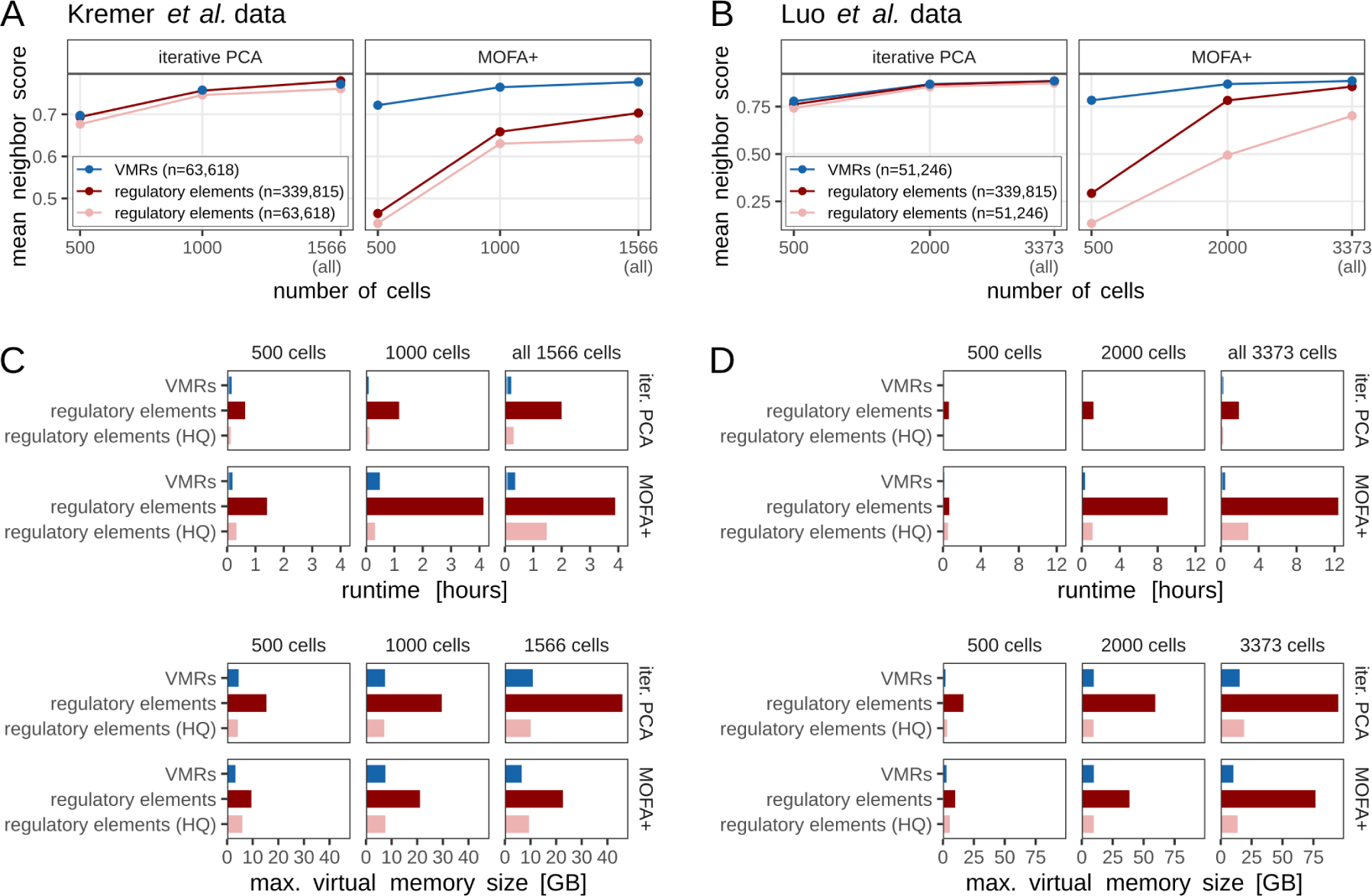
VMRs are more informative than an equal number of regulatory regions with high sequencing coverage. **(A-B)** Mean neighbor score obtained when quantifying CpG methylation at VMRs, all ENCODE regulatory regions, or a subset of regulatory regions for the data sets of Kremer et al. (2022) and Luo et al. (2017). The subset of high-quality (HQ) regulatory regions was selected in such a way that it matches the number of VMRs and contains those regulatory regions with the highest coverage. **(C-D)** Runtime (top) and RAM usage (bottom) of the two dimensionality reduction techniques shown in A and B.

**Figure S4:**
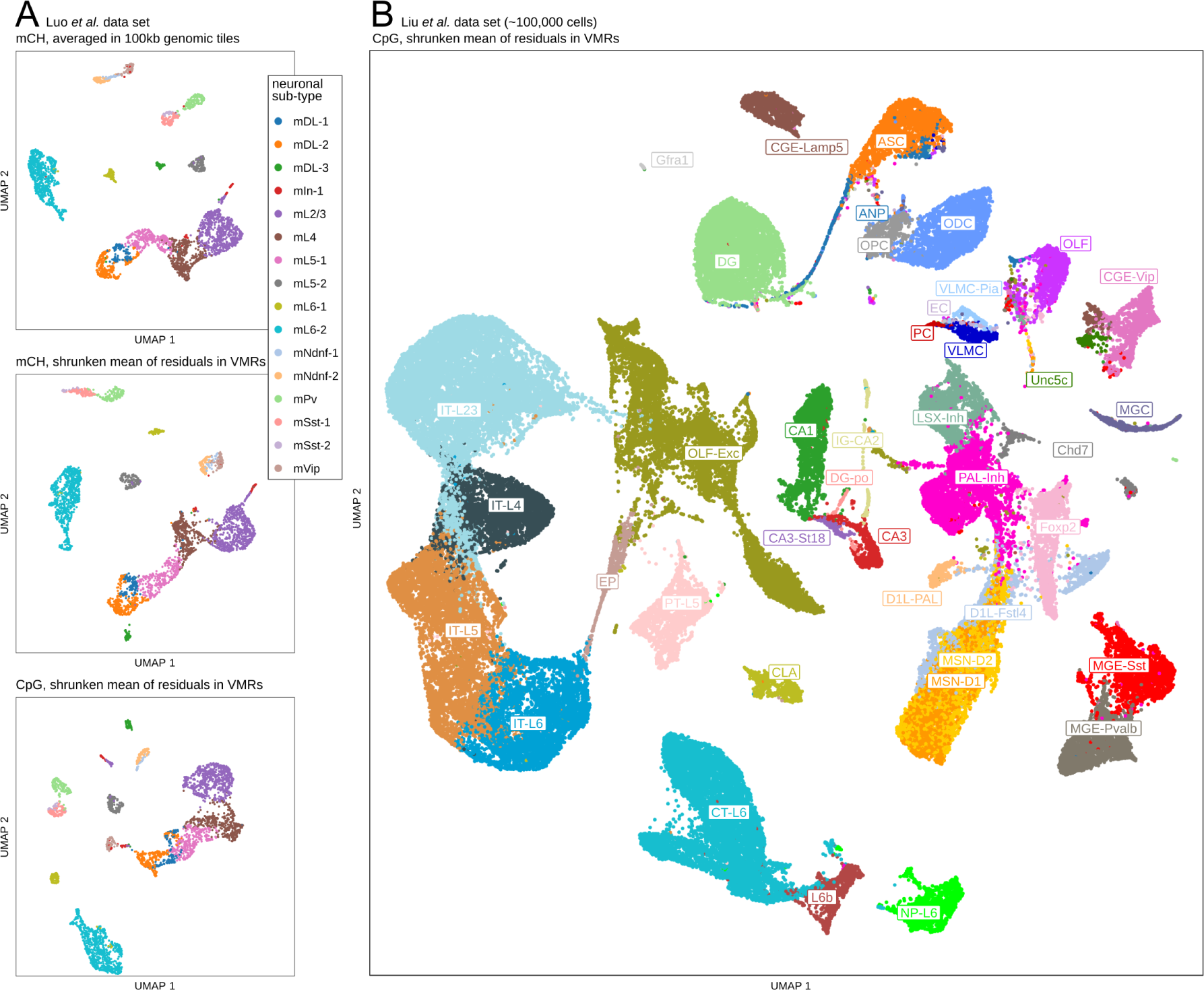
*MethSCAn* performs well on CH-methylation data and on extremely large data sets. **(A)** UMAPs obtained when analysing data from Luo et al. (2017) with three different strategies for producing a methylation matrix: top: averaging CH-methylation (mCH) in 100 kb genomic tiles (as done in Luo et al., 2017, to obtain the depicted cell type labels); center: using the shrunken mean of residuals to quantify mCH in mCH-VMRs; bottom: shrunken mean of residuals to quantify CpG methylation in CpG-VMRs. **(B)** UMAP obtained when analysing 100 350 neural single-cell methylomes (Liu et al., 2021) with the default *MethSCAn* workflow. The depicted cell labels were reported by Liu et al. (2021). Analysing this large data set takes approximately a week on a computer with 256 GB RAM and 48 CPUs, with almost all of this time (152 hours) spent by the prepare step, which is run only once per data set. The speed of this step can be increased considerably by processing chromosomes in parallel, which reduced the total runtime to approximately two days in our case.

